# Isoform-specific aPKC renders primary cilia dispensable for Hedgehog signaling and basal cell carcinoma growth

**DOI:** 10.1101/2020.06.08.137216

**Authors:** Tuyen T. L. Nguyen, Kirsten N. Wong, Ung Seop Jeon, Vama Jhumkhawala, Daniel Luy, Kevin C. Tan, Ishini Wickramatunga, Vinay Kumar, Whitney E. England, Linda T. Doan, Robert C. Spitale, Scott X. Atwood

**Affiliations:** Department of Developmental and Cell Biology, University of California, Irvine, Irvine, CA 92697, USA; NSF-Simons Center for Multiscale Cell Fate Research, University of California, Irvine, Irvine, CA 92697, USA; Department of Pharmaceutical Sciences, University of California, Irvine, Irvine, CA 92697, USA; Department of Dermatology, University of California, Irvine, Irvine, CA 92697, USA; Center for Complex Biological Systems, Chao Family Comprehensive Cancer Center, University of California, Irvine, Irvine, CA, 92697, USA

## Abstract

Primary cilia loss is a common feature of advanced cancers. While primary cilia are necessary to initiate Hedgehog (HH)-driven cancers, how HH pathway activity is maintained in advanced cancers devoid of primary cilia is unclear. Here, we find that HH-driven basal cell carcinoma (BCC) accumulates mutations in Alström and Usher syndrome genes. Loss of Alström and Usher syndrome gene expression, which are common underlying causes of deafness and blindness, suppresses primary ciliogenesis and HH signaling but enhances expression of atypical protein kinase C iota/lambda (aPKC), a GLI1 kinase necessary for advanced BCC growth. We show that aPKC expression is inversely correlated with primary ciliogenesis and that superficial BCCs display less primary cilia and higher aPKC expression, with the opposite trend in nodular BCC subtypes. Surprisingly, a constitutively active isoform of aPKC but not full-length protein drives HH pathway activity. Overexpression of the constitutively active aPKC variant can maintain HH pathway activity and tumor growth in the absence of primary cilia. Our results suggest tumors enhance isoform-specific expression of aPKC to prevent mutation-induced cessation of tumor growth.

## Introduction

The primary cilium is a non-motile, centrosome-based, microtubule signaling and sensing organelle that emanates from most mammalian cells after exiting from mitosis (Satir et al. 2010). Primary cilia house proteins that transduce many signals, including the Hedgehog (HH) signaling pathway, which are required during development and homeostasis. Alterations in signaling or structural components have been implicated in multiple ciliopathies and cancer (Hildebrandt et al. 2011), such as basal cell carcinoma (BCC). BCCs are locally invasive epithelial tumors that affect over 4 million people in the United States annually (Nguyen et al. 2019). BCCs require high sustained HH signaling for growth but some tumors paradoxically lose primary cilia during later stages while maintaining HH pathway activation (Atwood et al. 2015; Kuonen et al. 2019). Loss of primary cilia seems to be a general process in late-stage cancers with unclear significance and can be observed in advanced or metastatic breast cancer (Yuan et al. 2010), renal cancer (Schraml et al. 2009), prostate cancer (Hassounah et al. 2013), pancreatic cancer (Seeley et al. 2009), and melanoma (Kim et al. 2011).

In vertebrates, the HH signaling pathway is essential for embryonic development (Varjosalo and Taipale 2008) and regulates proliferation, migration, angiogenesis, and stem cell regeneration in adult tissues (Zheng et al. 2010). Vertebrate HH signals through the primary cilium by shutting down the cholesterol transporter PTCH1, allowing cholesterol accumulation and activation of the G-protein coupled receptor SMO and subsequent activation of the transcription factor GLI2, which transcribes HH target genes including the signal amplifier *GLI1* (Zhang et al. 2018). Inappropriate HH pathway activation drives BCC initiation and progression, mostly through mutations that inhibit PTCH1 (∼70%) or activate SMO (∼20%) (Bonilla et al. 2016). Loss of primary cilia typically leads to attenuation of HH signaling, however concomitant expression of GLI2 can accelerate BCC tumorigenesis (Han et al. 2009; Wong et al. 2009) in a process not well understood.

Atypical Protein Kinase C iota/lambda (aPKC or *PRKCI*) has been shown to be necessary to activate HH signaling in BCC and non-small cell lung cancer (Atwood et al. 2013; Yuan et al. 2007). Developmentally, aPKC plays a central role in determining cell polarity (Roignot et al. 2013), which is critical for actin-mediated regulation of primary ciliogenesis and axoneme length (Drummond et al. 2018), and can function as an oncogenic kinase where it phosphorylates the zinc finger domain of GLI1 to activate DNA binding and transcriptional activity (Atwood et al. 2013). Inhibition of aPKC using a peptide inhibitor shuts down HH signaling and tumor growth in a murine model of BCC (Atwood et al. 2013). Recently, we observed an increase in patient-derived mutations within cilia-associated genes in advanced skin cancer (Kuonen et al. 2019), where a subset of BCCs converted to squamous cell carcinoma (SCC) by shutting down the HH pathway and increasing RAS/MAPK signaling (Kuonen et al. 2019). These converted SCCs represent a small percentage of the total advanced BCCs which largely maintain HH pathway activation (Atwood et al. 2015). How HH signaling is maintained in advanced tumor growth when primary cilia are disrupted and the role aPKC plays in this process is unclear.

In this study, we find that loss of highly mutated cilia-associated genes disrupts primary cilia and attenuates HH pathway activation. Loss of primary cilia results in higher aPKC expression, which normally serves to restrict ciliogenesis. Analogously, superficial BCC tumors show a decrease in primary cilia and a concomitant increase in aPKC expression, which is inversely correlated to nodular BCC subtype, suggesting the mutual antagonism between aPKC and primary cilia may be functionally relevant in different BCC subtypes. Finally, expression of an aPKC isoform that only contains the kinase domain (aPKC-K), promotes HH signaling and tumor growth in the absence of primary cilia, indicating that aPKC-K may be the oncogenic driver of advanced BCC.

## Results

### Alström and Usher syndrome gene disruption suppresses primary cilia and HH signaling

We previously reported that 303 cilia-specific genes comprising the SYSCILIA list (van Dam et al. 2013) had significantly lower mutation rates in human BCCs compared to genome-wide mutation rates (Bonilla et al. 2016; Kuonen et al. 2019). However, many advanced BCC tumors displayed clonal loss of primary cilia (Kuonen et al. 2019), suggesting that some of the cilia-specific genes may be mutated more than normal. When we analyzed 161 cilia-specific genes that contained at least one mutation in human BCCs (Atwood et al. 2015), we found a subset of highly mutated genes in advanced BCCs compared to normal aged skin **(Figure 1A, Data S1)**. *PTCH1*, a negative suppressor of the HH pathway, is highly mutated as expected. *ALMS1*, which encodes a centrosomal protein mutated in Alström syndrome (Hearn 2019), is the most highly mutated gene in advanced BCC. Interestingly, four Usher syndrome-related genes, which are a major cause of deaf-blindness in humans (Géléoc and El-Amraoui 2020), are overrepresented in the top 9 most mutated genes. Not known to cause defects in primary cilia or HH signaling, Usher Type 1 proteins (CDH23, MYO7A, PCDH15, USH1C, and USH1G) connect the apical tips of hair bundle stereocilia to each other whereas Usher Type 2 proteins (GPR98, PDZD7, USH2A, and WHRN) connect the basal portion of these stereocilia. To determine differences in the mutation rate of Usher syndrome genes to the rest of the genome, we utilized both Gorlin’s syndrome patient samples (Chiang et al. 2018) and advanced human BCCs. When we compare Usher Type 1 and 2 genes, we observe significant increases in mutation rate from Gorlin’s and advanced BCCs, whereas normal aged skin or patient-matched blood samples do not change or decrease, respectively **(Figure 1B)**. These data suggest that highly mutated ciliopathy-associated genes from Alström and Usher syndromes may be the underlying cause of primary cilia loss in advanced cancer.

**Figure 1.**
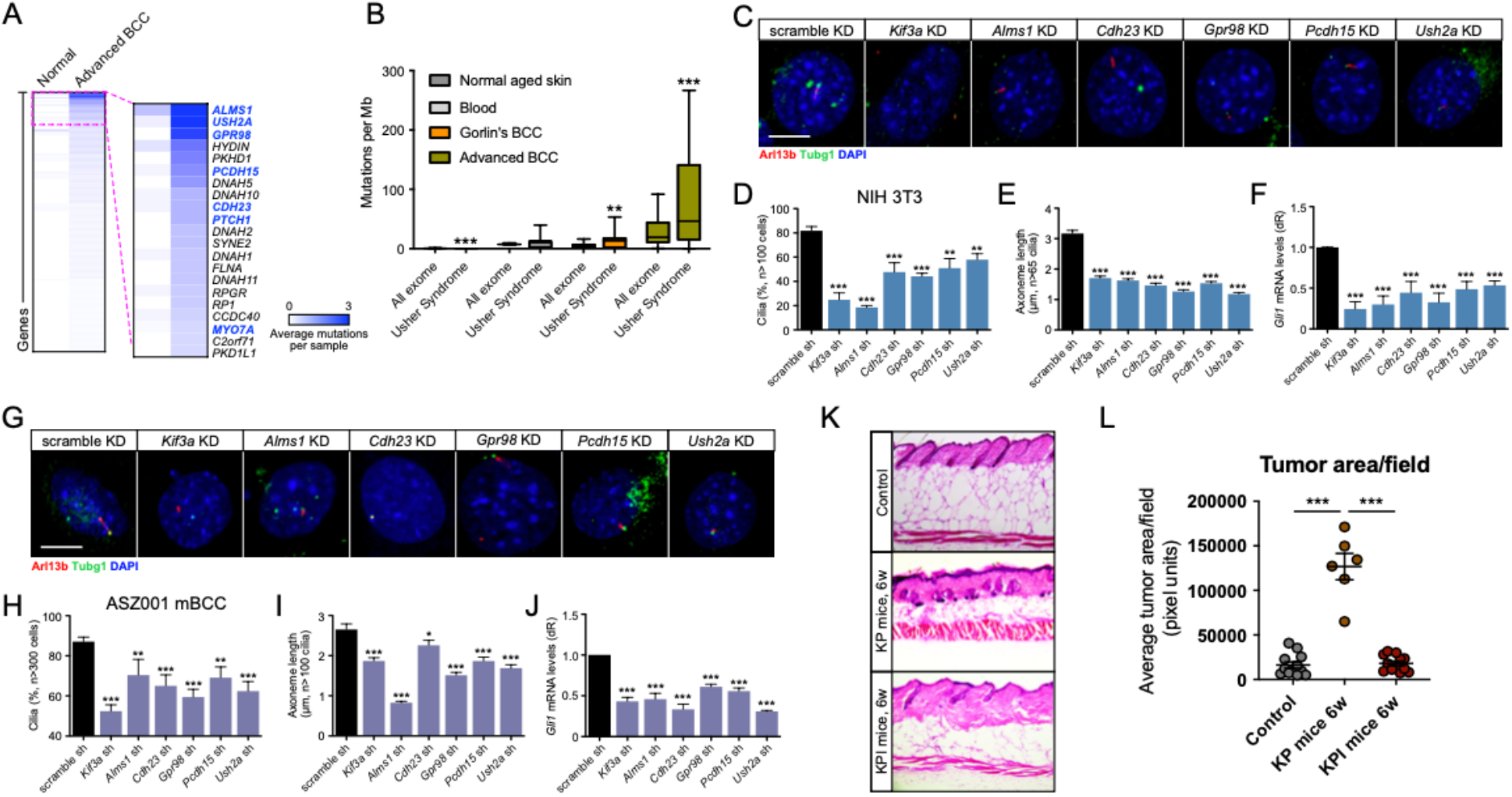
Alström and Usher syndrome genes are necessary to maintain primary ciliogenesis and HH signaling. **(A)** 161 mutated ciliary genes from exome sequencing of normal aged human skin (n=7) and advanced BCCs (n=15). **(B)** Mutations per megabase (Mb) of Usher syndrome genes versus background in blood (n=25), normal aged skin (n=7), Gorlin syndrome BCCs (n=21), and advanced BCCs (n=24). **(C-J)** Representative immunofluorescent images **(C, G)**, percentage (%) of ciliated cells **(D, H)**, average axoneme length **(E, I)**, and *Gli1* mRNA levels **(F, J)** of shRNA (sh)-mediated knockdown (KD) in NIH3T3 or ASZ001 cells and stained for the indicated markers. n ≥ 3. dR, delta reporter signal. Scale bar 10 µm. **(K-L)** H&E staining **(K)** and quantification of average tumor area per image field **(L)** of control (non-induced), *Krt14-*CreERT2*; Ptch1^fl/fl^* (KP), or *Krt14-*CreERT2*; Ptch1^fl/fl^*; *Ift88^fl/fl^* (KPI) mouse dorsal back skin. n ≥ 4 (KP n=2), 3 representative image fields per mouse. Error bars represent SEM; significance was determined by unpaired two-tailed *t* test (*, p<0.05; **, p<0.01; ***, p<0.001).

Despite the clonal loss of primary cilia in many advanced human BCCs, many of these tumors still maintain HH target gene induction (Atwood et al. 2015; Kuonen et al. 2019). To explore the possibility that disrupting expression of *Alms1* and Usher syndrome genes positively influences primary ciliogenesis and HH signaling, we used shRNA to knock down (KD) expression of each gene in HH-responsive NIH3T3 murine fibroblasts. We used *Kif3a* KD as a positive control as this gene is known to be required for primary ciliogenesis and HH signaling (Huangfu et al. 2003; Marszalek et al. 1999; Takeda et al. 1999). As expected, *Kif3a* KD resulted in loss of primary cilia, a reduction in axoneme length, and suppressed HH signaling compared to a scramble shRNA control **(Figures 1C-F, S1A)**. *Alms1* KD phenocopied *Kif3a* KD **(Figures 1C-F, S1A)**. Alms1 is a basal body protein that may control trafficking of vesicles into and out of the primary cilium and the *Alms1^-/-^* mice show many of the hallmarks of classical ciliopathies where cilia function is disrupted (Collin et al. 2005). Despite not being previously linked to primary cilia or HH signaling, KD of Usher Type 1 genes *Cdh23* and *Pcdh15* or Usher Type 2 genes *Gpr98* and *Ush2a* also resulted in primary cilia loss, reduced axoneme length, and diminished HH signaling compared to scrambled shRNA control **(Figures 1C-F, S1A)**. To rule out a cell-type specific role in primary ciliogenesis and HH signaling, we used ASZ001 murine BCC cells to KD gene expression and found comparable reductions in primary cilia frequency, axoneme length, and HH signaling **(Figures 1G-J, S1B)**. To verify that loss of primary cilia suppresses BCC tumor formation, we compared *Krt14-*CreERT2; *Ptch1^fl/fl^* (KP) mice, which initiates BCC growth after tamoxifen injection, to cilia-deficient *Krt14-*CreERT2; *Ptch1^fl/fl^; Ift88^fl/fl^* (KPI) mice and found significant suppression of tumor growth **(Figure 1K-L)**, in accordance with previous results (Wong et al. 2009). These data suggest that disruption of primary cilia-related genes does not allow for the maintenance of HH signaling and may need additional alterations to drive tumor growth.

### Primary cilia inhibit aPKC expression

As disruption of primary cilia does not maintain HH signaling or tumor growth by itself, we suspected that advanced tumors may activate other signaling components that drive HH signaling but are normally suppressed by primary cilia. To explore this idea further, we narrowed our focus to activators of the GLI transcription factors as previous work indicated that primary cilia loss combined with GLI2 expression, but not activation of SMO, accelerated BCC and medulloblastoma tumor growth (Han et al. 2009; Wong et al. 2009). As GLI1/GLI2 somatic copy number alterations are seen in only 8% of human BCCs and would not apply to most advanced tumors (Bonilla et al. 2016), we first explored whether the GLI1 activating kinase aPKC is regulated by primary cilia as aPKC is highly expressed in advanced BCC tumors (Atwood et al. 2013). When primary cilia are disrupted by *Kif3a*-, *Gpr98-*, or *Cdh23*-mediated KD, aPKC protein expression was significantly increased (**Figure 2A-B**). This result was unexpected as aPKC is a target gene of GLI1 and one would expect a reduction in expression if HH signaling was disrupted. However, this effect also occurred *in vivo* as close examination of allografted murine BCCs generated from *Krt14*-CreERT2; *Ptch1*^+/-^; *Trp53^fl/fl^* (KPT) mice revealed differential aPKC expression that anti-correlated with primary cilia within each sample. Higher aPKC expression coincided with lower ciliation and lower aPKC expression coincided with higher ciliation (**Figure 2C-D**), suggesting aPKC and primary cilia may mutually inhibit each other. In fact, overexpression of aPKC in NIH3T3 cells resulted in a significant decrease in primary ciliogenesis (**Figure S2A-B**).

**Figure 2.**
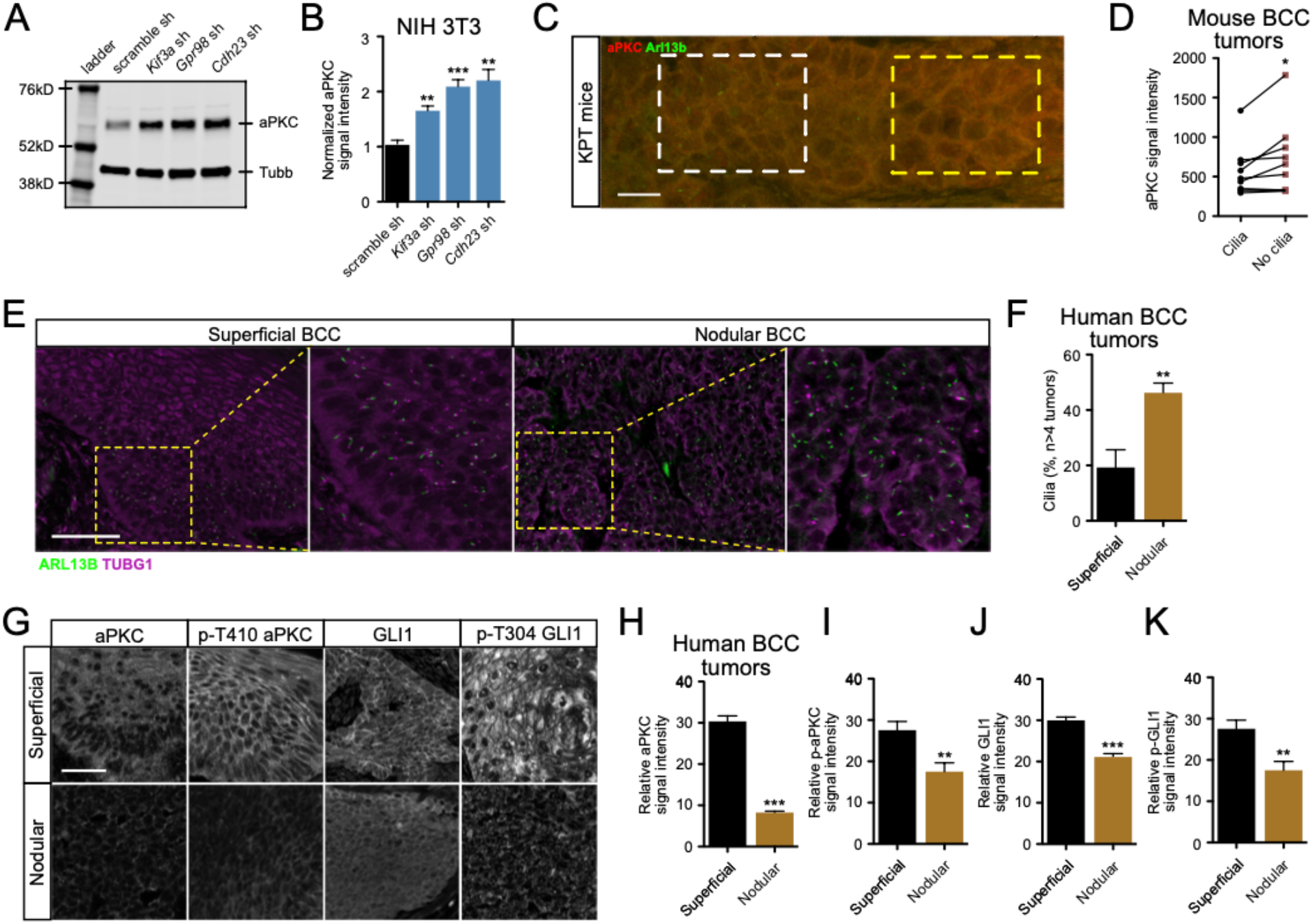
Primary cilia inhibit aPKC expression and are differentially regulated in BCC subtypes. **(A-B)** Western blot **(A)** and quantification **(B)** of NIH3T3 cells with the indicated shRNA-mediated KD. kD, kilodalton. n = 4. **(C-D)** Immunofluorescence **(C)** and quantification **(D)** of adjacent regions in *Krt14*-CreERT2*; Ptch1^+/-^; Trp53^fl/fl^* (KPT) irradiated mice. Scale bar, 10 µm. White dotted square: ciliated section. Yellow dotted square: section without cilia. n = 9. **(E)** Superficial or nodular human BCCs stained with the indicated markers. Scale bar, 50 µm. **(F)** Quantification of ciliated cells. n ≥ 4 tumors. **(G)** Superficial or nodular human BCCs stained with the indicated antibodies. Scale bar, 50 µm. **(H-K)** Quantification of tumor immunofluorescence. N ≥ 4 tumors. Error bars represent SEM; significance was determined by unpaired two-tailed *t* test (*, p<0.05; **, p<0.01; ***, p<0.001).

To better define the aPKC and primary cilia relationship *in vivo*, we quantified primary cilia and aPKC expression in nodular and superficial human BCC subtypes. Nodular BCCs are thought to arise from hair follicles (Grachtchouk et al. 2011), whereas superficial BCCs arise from the interfollicular epidermis (Tan et al. 2018). Our results showed significantly higher primary ciliogenesis in nodular BCCs compared to superficial tumors (**Figure 2E-F**). At the same time, the average expression of total aPKC and activated aPKC (phosphorylated at T410) proteins were significantly lower in nodular BCCs with more primary cilia than in superficial tumors with less primary cilia (**Figure 2G-I**). Similarly, total GLI1 and activated GLI1 (phosphorylated at T304 by aPKC) were also significantly reduced in nodular BCCs **(Figure 2G, J-K)**, suggesting that superficial tumors may require more HH activation to grow than nodular tumors.

### Constitutively active aPKC isoform promotes HH signaling

To verify that aPKC is highly active in human BCCs, we took a bioinformatics approach and inferred an aPKC-specific gene signature list from RNA-seq analysis of BCC cells treated with aPKC or SMO antagonist (Atwood et al., 2013). The vast majority of differentially expressed transcripts are common between the two inhibitors, with 5% of transcripts unique to aPKC antagonist and 10% unique to SMO antagonist **(Data S2)**. We compared each gene list to RNA-seq datasets of 14 tumor-normal pairs of advanced BCCs (Atwood et al. 2015) and found 34 aPKC-specific, 76 SMO-specific, and 736 commonly shared transcripts that were differentially expressed by two-fold or more **(Figure 3A, Data S2)**. Two-thirds of each gene list showed significant increases in gene expression compared to the matched normal sample. Upregulated genes in the aPKC-specific response correspond to cell cycle, cancer-related pathways, and the ciliary landscape according to WikiPathways analysis (Slenter et al. 2018) **(Figure 3B, Data S3)**. Notable genes include the MCM genes and their downstream target UBEC2, which are involved in the transcriptional control of ciliogenesis and centrosome amplification (Casar Tena et al. 2019).

**Figure 3.**
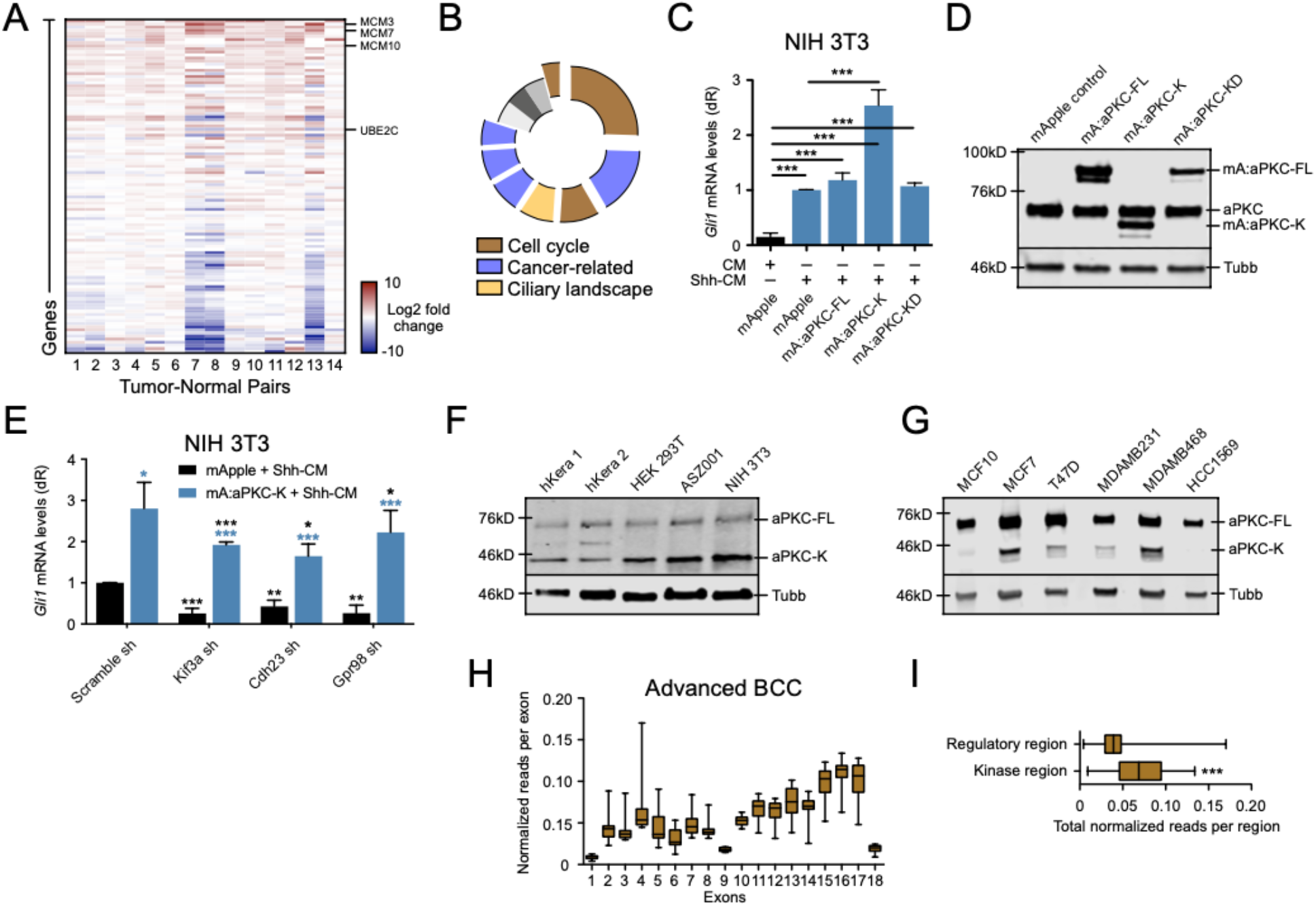
Constitutively active aPKC isoform promotes HH signaling. **(A)** aPKC signature gene expression in 14 advanced human BCC-normal pairs. **(B)** Wikipathways analysis of upregulated aPKC-dependent genes. **(C)** *Gli1* mRNA levels of NIH3T3 cells overexpressing the indicated constructs with conditioned media (CM) or Shh-CM. n ≥ 5. dR, delta reporter signal. mA, mApple. FL, full-length. K, kinase. KD, kinase dead. **(D)** Western blots of NIH3T3 cells overexpressing the indicated aPKC constructs. Top blot: C-terminal aPKC antibody. kD, kilodalton. **(E)** *Gli1* mRNA levels of NIH 3T3 cells expressing mApple or mApple-aPKC-K containing the indicated shRNAs (sh). n ≥ 3. Black stars: p values compared to mApple scramble sh DMSO control. Blue stars: p values compared to internal mApple controls. **(F-G)** Western blots of the indicated cell lines. hKera, primary human keratinocyte. **(H)** Normalized reads per exon of the *PRKCI* gene from 14 advanced human BCCs. Data normalized to read length. **(I)** Total normalized reads per region of the *PRKCI* gene. For box and whisker plots: box represents 25^th^ to 75^th^ percentiles; whiskers represent minimum and maximum data points; bar represents mean. For bar plot: error bars represent SEM. Significance was determined by unpaired two-tailed *t* test (*, p<0.05; **, p<0.01; ***, p<0.001).

As aPKC’s gain-of-function role in HH signaling has not been tested, we overexpressed an mApple-tagged full-length wild-type isoform of aPKC (aPKC-FL) or an mApple-tagged kinase-dead variant (aPKC-KD) in NIH3T3 cells but did not observe changes in HH signaling in the presence of Shh-CM as measured by *Gli1* mRNA levels, a HH target gene (**Figure 3C-D**). Since the loss of aPKC plays a significant role in suppressing HH pathway activity (Atwood et al. 2013), we hypothesized that subcellular location or an alternatively spliced variant may be responsible for promoting HH activity. Interestingly, an mApple-tagged constitutively active isoform of aPKC, where the regulatory amino-terminal domain is truncated (aPKC-K; amino acids 235-587), showed more nuclear localization compared to full-length aPKC and significantly increased HH signaling **(Figures 3C-D, S2C)**. To characterize the role of aPKC-K overexpression when primary cilia are disrupted, we treated NIH3T3 cells stably expressing aPKC-K or a vector control with shRNA-mediated KD of relevant cilia genes. Scramble control cells showed a significant increase in *Gli1* mRNA levels upon aPKC-K overexpression as expected (**Figure 3E**). Intriguingly, when we disrupted primary cilia through shRNA-mediated KD of *Kif3a*, *Cdh23*, or *Gpr98*, we found that all conditions showed an expected reduction in *Gli1* mRNA levels in control cells but an increase in *Gli1* levels in the presence of aPKC-K **(Figure 3E)**. In fact, aPKC-K protein significantly induced *Gli1* expression at a higher level than intact scramble control cells, suggesting that aPKC-K not only maintains HH pathway activity when primary cilia are disrupted but can elevate expression above baseline levels.

The aPKC-K isoform, otherwise known as PKMζ, has been shown to play an important role in long-term memory potentiation (Tsokas et al. 2016). To determine whether this isoform is present outside of brain tissue, we performed western blots of a variety of cell types and observed both endogenous aPKC-FL and aPKC-K isoform expression in primary human keratinocytes, human embryonic kidney 293T cells, mouse NIH3T3 fibroblasts, and mouse ASZ001 BCC cells using an antibody that recognizes the kinase domain of aPKC **(Figure 3F)**. We also observed both endogenous aPKC-FL and aPKC-K isoform expression in western blots of normal human breast epithelial cell line MCF10 and multiple human breast cancer cell lines **(Figure 3G)**. In addition, both endogenous bands are reduced upon *aPKC* KD **(Figure S2D)** and are elevated upon *Kif3a* KD **(Figure S2E)**. Finally, when we quantify reads per exon within the *PRKCI* gene from bulk-level RNA-seq of advanced human BCCs (Atwood et al. 2015), we observed more reads within exons 9-18 (corresponding to aPKC-K) than within exons 1-8 (corresponding to the regulatory region) **(Figure 3H-I)**. These data indicate that the aPKC-K isoform is widely expressed in normal and tumor contexts and the aPKC-K isoform can drive HH pathway activity.

### aPKC-K drives *in vivo* HH signaling and tumor growth in the absence of primary cilia

To determine whether aPKC-K can drive *in vivo* tumor growth, we generated a conditional aPKC-K expressing mouse under the control of Cre recombinase (aPKC-K^cond^). A targeting cassette encoding aPKC-K with an amino-terminal dTomato tag was used to target the Rosa26 locus 3’ to a LoxP-flanked polyadenylation stop sequence cassette in a strategy similar to transgenic SmoM2 animals (Jeong et al. 2004). When we crossed aPKC-K^cond^ to KPI mice (KPIA mice: *Krt14-* CreERT2; *Ptch1^fl/fl^; Ift88^fl/fl^*; aPKC-K^cond^), we observed significant tumor growth at 6 and 12 weeks post tamoxifen injection **(Figure 4A-B)**. Tumors grew from Krt14+ tissues in the epidermis and upper hair follicles. Primary cilia were significantly reduced in KPI and KPIA mice compared to KP mice with axonemes of remaining cilia showing significant reduction in length in both the hair follicle-derived tumors **(Figure 4C-E)** and epidermis and epidermal-derived tumors **(Figure 4F-H)**, suggesting that aPKC-K does not rescue primary cilia. Consistent with the mutual antagonism between aPKC and primary cilia in observed in cell culture **(Figures 2A-B, S2A-B)**, expression of aPKC-K in KPIA mice further decreased ciliated cells compared to KPI mice **(Figure 4G)**. Gli1 protein levels were significantly elevated in KPIA mice compared to KP mice **(Figure 4I-J)**, indicating that aPKC-K expression can drive *in vivo* HH signaling and tumor growth in the absence of primary cilia.

**Figure 4.**
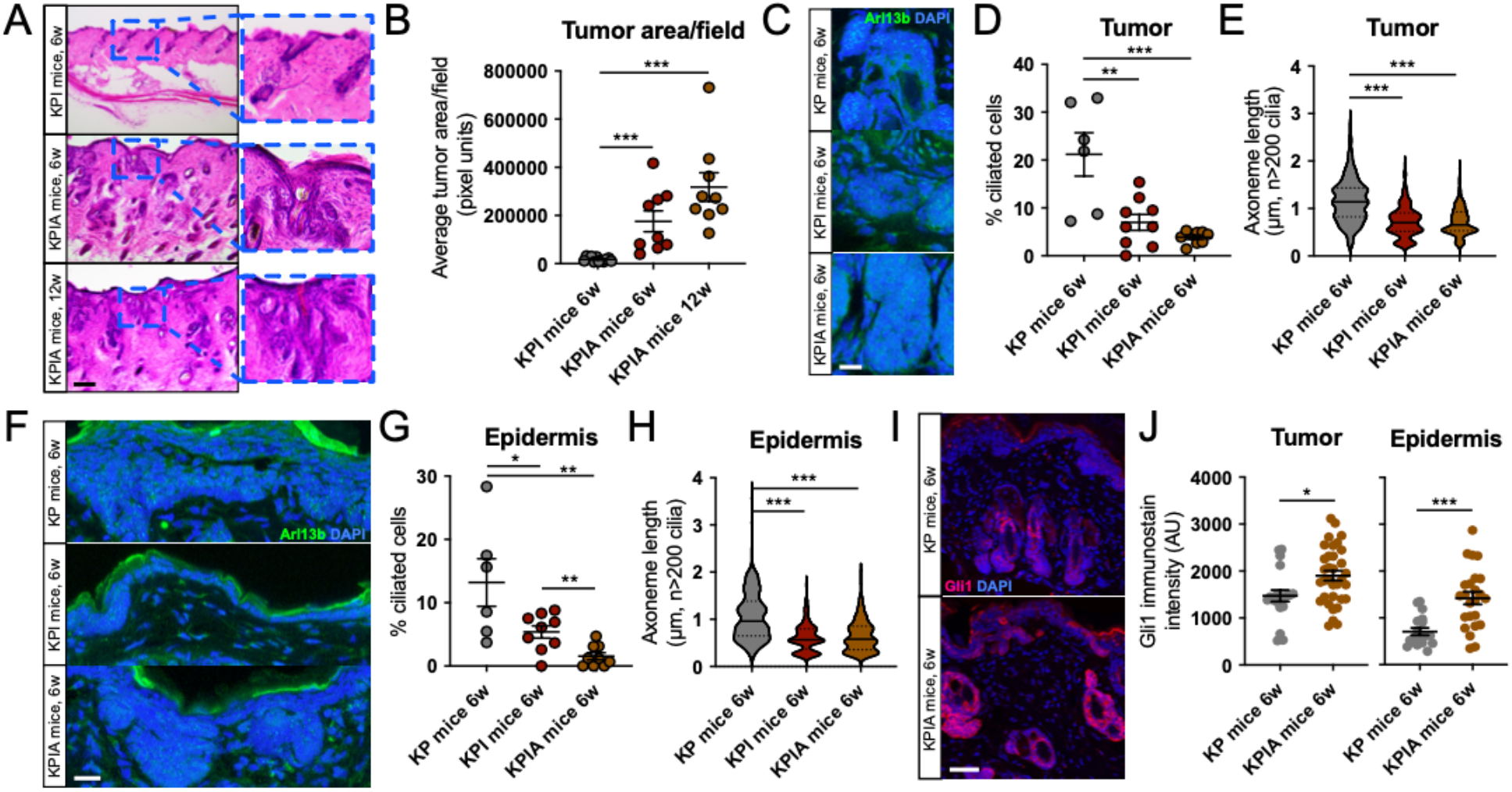
aPKC-K isoform promotes HH signaling and tumor growth in the absence of primary cilia. **(A-B)** H&E staining **(A)** and quantification of average tumor area per image field **(B)** of *Krt14-*CreERT2*; Ptch1^fl/fl^*; *Ift88^fl/fl^* (KPI) or *Krt14-*CreERT2*; Ptch1^fl/fl^*; *Ift88^fl/fl^; aPKC^cond^* (KPIA) mouse dorsal back skin. n ≥ 3, 3 representative image fields per mouse. 6w, 6 weeks post induction. 12w, 12 weeks post induction. Scale bar, 100 µm. **(C-H)** Immunofluorescence **(C, F)** and quantification of primary cilia-containing cells **(D, G)** or individual axoneme length **(E, H)** in hair follicle-derived tumors or epidermis/epidermal tumors (3 representative image fields per mouse) of dorsal back skin from *Krt14-*CreERT2*; Ptch1^fl/fl^* (KP) mice or the indicated genotypes using Arl13B (cilia) and DAPI. n ≥ 3 (KP n=2). Scale bar, 20 µm. **(I-J)** Immunofluorescence **(I)** and quantification **(J)** of Gli1 immunostain intensity in hair follicle-derived tumors or epidermis/epidermal tumors of the indicated genotypes from ≥ 8 representative regions per mouse (KP n=2, KPIA n=3). Scale bar, 20 µm. AU, arbitrary units. Error bars represent SEM; significance was determined by unpaired two-tailed *t* test (*, p<0.05; **, p<0.01; ***, p<0.001).

## DISCUSSION

The HH signaling pathway is the driving force in BCC formation and maintenance (Epstein 2008), which is heavily dependent on intact primary cilia (Satir et al. 2010). Yet, HH signaling remains elevated in many advanced BCCs where primary cilia are significantly disrupted. How Alström and Usher syndrome genes regulate primary ciliogenesis and HH signaling is unclear. ALMS1 has been identified through AP-MS studies as a possible binding partner of GPRASP2, a GPCR-associated sorting protein that has been implicated in SMO ciliary translocation (Jung et al. 2016). PCDH15 localization to kinocilia is dependent on FGFR1 activity (Honda et al. 2018), an interesting connection given that FGFR regulates GLI1 expression downstream of SMO in medulloblastomas (Neve et al. 2019). In addition, *USH2A* and *PCDH15* can both bind directly to GBX2 (Roelseler et al. 2012), a HH-responsive protein in thalamic differentiation (Szabó et al. 2009). Moreover, CDH23 is a Dachsous-like protein that may regulate HH signaling given that Dally and Dally-like, the downstream targets of tumor suppressor Dachsous, have been shown to modulate the HH signaling pathway in *Drosophila* (Williams et al. 2010).

Ciliogenesis is essential to maintain cellular homeostasis, specifically to facilitate proper signaling of the HH, WNT, NOTCH, Hippo, PDGFRα, FGF, and other GPCR pathways in certain contexts (Wheway et al. 2018). Many of these pathways facilitate tumor growth. So primary cilia loss in advanced tumors is unexpected from a mechanistic view and how it benefits tumor growth is unclear. Generating primary cilia during the G1 portion of the cell cycle is energetically expensive and disruption of this process in certain genetic backgrounds may allow for faster cycling of tumor cells, offsetting loss of the cilia-mediated oncogenic signal (Liu et al. 2018). This would have to occur after tumor initiation, as our data and others indicate that simultaneous loss of primary cilia and the oncogenic event suppresses tumor growth (Wong et al. 2009). A likely order of events in our context would be elevated aPKC expression given that aPKC is a HH target gene (Atwood et al. 2013), accumulation of cilia-related gene mutations, and subsequent loss of primary cilia. Alternatively, certain oncogenic signals may be restrained by primary cilia-mediated signals. aPKC may serve as this type of oncogenic signal as primary cilia restrain aPKC expression, which is relieved upon cilia loss.

aPKC is overexpressed in many cancer models including ovarian (Eder et al. 2005), lung (Regala et al. 2005), brain (Baldwin et al. 2008), esophagus (Yang et al. 2008), breast (Kojima et al. 2008), and pancreatic (Evans et al. 2003) cancers, where it is often tied to its cell polarity functions. Here we show that the constitutively active aPKC-K isoform activates HH signaling and likely serves as an oncogene in BCC. Our data are consistent with previous findings that aPKC-K overexpression, but not full-length aPKC, leads to the formation of large neuroblast tumors in *Drosophila* larval brains (Lee et al. 2006). aPKC-K does not polarize like its full-length counterpart in neuroblasts, suggesting that aPKC-K’s role in HH target gene induction may be independent of polarity. The aPKC-K isoform is also required for mouse brain development (Hirai et al. 2003) and long-term memory potentiation (Tsokas et al. 2016), indicating critical roles for this isoform in the brain. We demonstrate aPKC-K expression in mammalian keratinocytes, embryonic kidney cells, normal and cancerous breast cells, fibroblasts, and BCC cells, suggesting that aPKC-K may also have essential roles in these tissues. Together, these findings suggest broad implications for isoform-specific expression of aPKC and reveal a role for its enhancement by tumors to prevent mutation-induced cessation of tumor growth.

## MATERIALS AND METHODS

### Data Availability Statement

Datasets related to this article can be found at https://www.ncbi.nlm.nih.gov/geo/query/acc.cgi?acc=GSE58377, hosted at GEO with reference number GSE58377 (Atwood et al. 2015) and National Institutes of Health Sequence Read Archive SAMN07507265−SAMN07507288 (Chiang et al. 2018).

### Generation of stable cell lines

PiggyBac transposons containing sequences for full-length wildtype *PRKCI* (aPKC-FL), full-length K274W *PRKCI* (kinase dead, aPKC-KD), or *PRKCI* 235-587 (constitutively active, aPKC-K) were co-transfected with piggyBac transposase using Lipofectamine 3000 reagent (L3000150, LifeTechnologies) per manufacturer’s protocols. Transfected cells were selected in 500µg/mL Geneticin G-418 (50841720, Fisher Scientific) for 1 additional passage after all non-transfected control cells died. Stable cells were then maintained in regular culture media for at least 3 additional passages prior to experiments.

### Lentiviral knockdown

Lentiviral pLKO.1 vector (Open Biosystems) containing shRNAs to *Prkci* (5′-CCAGAC AGAAAG CAGGTT GTT-3′; Atwood et al. 2013), *Kif3a* (5′-TCCGCC AGTTTC AGAAAG AAA-3′); or *pGIPZ* ires GFP vector containing shRNAs to *Alms1* (5’-TAGAAG TTAGTT TGTCCTG GC-3’), *Cdh23* (5’-ATTGAT GACGAT CTTCAC GCG-3’), *Grp98* (5’-TATAAA GTCTAT CATTGG TGG-3’), *Pcdh15* (5’-AATAGG GTTCAG TTCTTC CGG-3’), and *Ush2a* (5’-TTGATG ATAATG TGTCGC G-3’) were used. Lentiviral infection was performed on ASZ001 and NIH3T3 cells and assayed between 4 and 6 days depending on the efficiency of knockdown, as determined by mRNA levels using RT-qPCR.

### Generation of transgenic animals

To generate the aPKC-K knock-in mouse, we contracted Cyagen Biosciences Inc. to use NCBI reference sequence: NM_008857.3 on chromosome 3 to amplify the kinase domain isoform of mouse *Prkci*. A “dTomato (without a stop codon)-2A-*Prkci* kinase domain-polyA” cassette was cloned into intron 1 of ROSA26, with a CMV-LoxP-Stop-LoxP placed upstream such that expression of the cassette is dependent on expression of Cre recombinase in C57BL/6 ES cells. Neomycin was used for positive selection and DTA for negative selection, with the Neo cassette later removed. ES clones were confirmed via Southern Blot, microinjected into a blastocyst, followed by chimera production. Founders were confirmed as germline-transmitted via crossbreeding with wild-type mice. All transgenic mice used in this study were confirmed via genotyping.

## ACKNOWLEDGEMENTS

We wish to thank Dr. Mike Drummond for his assistance with troubleshooting in the design of aPKC constructs, Dr. Vaishali Jayashankar for provision of breast epithelial and cancer cell lines, as well as all members of the Atwood lab for their constant support. S.X.A. is supported by National Institutes of Health (NIH) grant R01CA237563 and American Cancer Society Research Scholar Award RSG-19-089-01-DDC. T.T.L.N. is supported by a National Science Foundation Graduate Research Fellowship Program Award and a Graduate Assistance in Areas of National Need Fellowship. K.N.W. is supported by NIH National Cancer Institute (NCI) training grant CA009054 and Simons Foundation grant 594598. The authors wish to acknowledge the support of the Chao Family Comprehensive Cancer Center Genomics High-Throughput Facility and Optical Biology Core Shared Resource, supported by the NIH NCI award P30CA062203 and the UCI Skin Biology Resource Center supported by the NIAMS award P30AR075047.

## CONTRIBUTIONS

S.X.A. and T.T.L.N. conceived the project; S.X.A. supervised research; T.T.L.N. and K.N.W. performed experiments; T.T.L.N., K.N.W., U.S.J., V.J., K.C.T., I.W., and S.X.A. quantified experimental data; V.K., W.E.E., and R.C.S. analyzed RNA-seq data; L.T.D. collected and annotated human clinical samples; S.X.A., T.T.L.N., and K.N.W. wrote the manuscript. All authors analyzed and discussed the results and commented on the manuscript.

## Supplemental Methods

### Ethics statements

Human clinical studies were approved by the Ethics Committee of the University of California, Irvine. All human studies were performed in strict adherence to the Institutional Review Board (IRB) guidelines of the University of California, Irvine (2009-7083).

### Human samples

Written informed consent was obtained for all archived human samples and was reviewed by the University of California Irvine IRB. Human normal epidermis and BCC samples were collected from UC Irvine Medical Center. Paraffinized samples were sectioned with a rotary microtome (Leica RM2155) at 7 μm for analysis. Samples were deparaffinized as described by Abcam and antigen retrieval was performed using a Tris-EDTA buffer (10 nM Tris base, 1 mM EDTA, 0.05% Tween 20, pH 9.0) at 100°C for 10 min.

### Mice

All mice were housed under standard conditions. Animal care was in compliance with the protocols approved by the Institutional Animal Care and Use Committee (IACUC) at the University of California, Irvine. Both male and female *Krt14-*CreERT2*; Ptch1^fl/fl^* (KP), *Krt14-*CreERT2*; Ptch1^fl/fl^*; *Ift88^fl/fl^* (KPI), and *Krt14-*CreERT2*; Ptch1^fl/fl^*; *Ift88^fl/fl^; aPKC^cond^*(KPIA) mice were injected with 100uL of 10 mg/mL tamoxifen (Sigma) intraperitoneally for three consecutive days at 6-10 weeks of age. Mice were euthanized and dorsal back skin samples were collected 6-12 weeks post-induction to assess tumor burden. Collected skin samples were fixed with 4% paraformaldehyde (PFA; Electron Microscopy Sciences) for 1 hour at room temperature, washed with DPBS (Life Technologies), immersed in 30% sucrose at 4°C overnight, and frozen in Tissue-Tek OCT Compound (Sakura Finetek). Samples were then cryo-sectioned using the CryoStar NX50 Cryostat (Thermo Fisher Scientific) at 10-20um for analysis. At least three mice were used for each time point or condition unless otherwise stated.

### RNA Sequencing Analysis

RNA-seq data were obtained from patient-matched advanced human BCC patients (Atwood et al. 2015). RNA-Seq data were aligned as previously described (Atwood et al. 2015). The NCBI Reference Sequence (RefSeq) databases were used as reference annotations to calculate values of reads per kilobase of transcript per million mapped reads for known transcripts (RPKM). RPKM values were then log2-transformed, and heat map analysis was used to visualize the differential gene expression. Pathway enrichment terms from RNA-seq data were obtained using Enrichr (Kuleshov et al. 2016). The RNA-seq data were quality trimmed using Trimmomatic (Bolger et al. 2014) and aligned to the ENSEMBL human reference transcriptome (GRCh38.93) modified to include the aPKC kinase domain (exons 9-18) with STAR (Dobin et al. 2013). Bedtools was used to extract the counts of read mapping to the exons of aPKC.

### Exome Sequencing Analysis

Exome-Seq data were aligned as previously described (Atwood et al. 2015; Chiang et al. 2018). Mutations per sample were calculated for the indicated genes and displayed using a heatmap. Mutations per megabase were calculated for each sample set with the sample-matched overall mutation rates (all exome) used as a control.

### Cell culture, drug treatments, and quantitative RT-PCR

ASZ001 cells (So et al. 2006) were grown in 154CF/PRF media kit (Life Technologies, M154CFPRF500) supplemented with 1% Penicillin/Streptomycin (Life Technologies, 15140122), 2% FBS (Life Technologies, 10437028) chelated overnight with Chelex® 100 Resin (Bio-Rad, 1422822), and 0.07mM CaCl2. NIH3T3 cells (ATCC, CRL 1658) were grown in DMEM media containing 10% FBS and 1% Penicillin/Streptomycin. MCF10A cells were grown in DMEM/F12 supplemented with 20% horse serum, EGF (20 ng/ml), hydrocortisone (0.5 mg/ml), cholera toxin (100 ng/ml) and insulin (10 μg/ml). MDA-MB-231, MDA-MB-468, and MCF7 cells were grown in DMEM with 10% FBS, insulin (10 μg/ml), and 1mM sodium pyruvate. MCF7 cells were additionally supplemented with 2mM L-Glutamine. T47D cells and HCC1569 cells were grown in RPMI supplemented with 10% FBS and insulin (10 μg/ml).

At confluence, cells were serum-starved +/− Shh-N ligand (1:100) for 24 hours. RNA was harvested with the Direct-zol RNA MiniPrep Plus kit (ZYMO Research, R2072). Quantitative RT-PCR was performed using the iTaq Universal SYBR Green 1-Step Kit (Bio-Rad, 1725151) on a StepOnePlus Real-time PCR system (Applied BioSystem) using primers for *Gli1* (forward: 5′-GCAGGTG TGAGGCC AGGTAG TGACGA TG-3′, reverse: 5′-CGCGGG CAGCAC TGAGGA CTTGTC-3′), *Gapdh* (forward: 5′-AATGAA TACGGC TACAGC AACAGG GTG-3′, reverse: 5′-AATTGT GAGGGA GATGCT CAGTGT TGGG-3′), *mApple* (forward: 5’-ACCTAC AAGGCC AAGAAG CC-3’, reverse: 5’-GCGTTC GTACTG TTCCAC GA-3’), *Kif3a* (forward: 5’-CAGACT GGGACA GGCAAG AC-3’, reverse: 5’-TTTCTT TTTACC TGCTTG GTCCCT-3’), *Alms1* (forward: 5’-TGTGCA GGAGTC TCATGG TT-3’, reverse: 5’-TCTCCC CAGGAG ATGATT GGT-3’), *Cdh23* (forward: 5’-ACTGGC TCACAG AGGAGA GT-3’, reverse: 5’-CAGAAG AGGCTT GCCCTG G-3’), *Grp98* (forward: 5’-AATAGT CTGCAG GGACTT TATGTT T-3’, reverse: 5’-CAGCTG GAGGTA TTCCGC TC-3’), *Pcdh23* (forward: 5’-TCTTGG CACAGG GACCTA CT-3’, reverse: 5’-CGTATT GCCAGT CATCGT CA-3’), *Ush2a* (forward: 5’-AACCCA TGTGGG ATTAGC CG-3’, reverse: 5’-GAGGTG GGTGTC GGTAAA GG-3’), and *Prkci* (forward: 5’-AAGGAA CGATTG GGTTGT CACCCT-3’, reverse: 5’-AAGGGT GGAACC ACCTGC TTTTGC T-3’). Fold change in mRNA expression of target genes was measured using ΔΔCt analysis with *Gapdh* as an internal control gene. Experiments were repeated at least three times and data was represented as the mean of triplicates ± SEM.

### Tumor Quantification

Skin sections (20μm) were stained with hematoxylin and eosin (H&E; Richard-Allan Scientific) per standardized protocol. Stained sections were imaged at 10x magnification using the AmScope microscope with the AmScope MU500B digital camera. Tumor area was measured using ImageJ. Tumors were assessed as the total tumor size per image field. Tumor area was quantitated from 3 representative fields per sample. The average was calculated for each condition.

### Immunofluorescence

Control, transfected, or virally infected cells were seeded in respective growth media. At confluence, the cells were serum starved for 24 hours, then fixed with 4% paraformaldehyde for 15 minutes, and blocked in PBS containing 1% normal horse serum and 0.1% Triton X-100 for 30 minutes. The following antibodies were used: rabbit anti-γ-tubulin (1:1000, SAB4503045; Sigma), mouse anti-ARL13B (1:1000, 75–287; Antibodies Inc.), rabbit anti-RFP (1:1000, RL600-401-379; Rockland), rabbit anti-aPKC (1:500, sc-216; Santa Cruz Biotechnology), rabbit anti-p-aPKC T410 (1:500, sc-12894; Santa Cruz Biotechnology), rabbit anti-GLI1 (1:500, AF3455; R&D Systems), and rabbit anti-P-T304 GLI1 (1:200) (Drummond et al. 2018). Secondary antibodies included Alexa Fluor 488, 546, and 647 (Jackson ImmunoResearch). Slides were mounted in Prolong Diamond Antifade Mountant with DAPI (P36962, Molecular Probes). Confocal images were acquired at room temperature on a Zeiss LSM700 laser scanning microscope with Plan-Apochromat 40× and 63× oil immersion objectives. For KP, KPI, and KPIA mouse tissues, confocal images were acquired on a Leica SP8 Confocal Microscope with 40x oil immersion objective. Fluorescent images were acquired at room temperature on an EVOS FL Color Imaging System with Plan Fluorite 40× objective. Images were arranged with ImageJ, Affinity Photo, and Affinity Designer.

### Western blotting

Cells or tissues were lysed with 2× SDS sample buffer (100 mM Tris HCl 6.8, 200 nM DTT, 4% SDS, 0.2% bromophenol blue, and 20% glycerol) and boiled at 95°C for 15 minutes. Samples were resolved on a 4–12% polyacrylamide gradient gel and transferred to nitrocellulose membrane by a wet transfer apparatus. Membranes were blocked with 5% milk in TBS with 0.05% Tween-20 before addition of antibodies: rabbit anti-aPKC (1:1000, sc-216; Santa Cruz Biotechnology), mouse anti–ß-Tubulin (1:5000, E7;DSHB), rabbit anti-RFP (1:1000, RL600-401-379; Rockland), and Alexa Fluor secondary antibodies. Membranes were imaged and quantified using the LI-COR Odyssey CLx imaging system with built-in Image Studio software.

### Statistics

Statistical analyses were done in GraphPad Prism using two-tailed *t* tests.

---

**Supplemental Figure 1.**
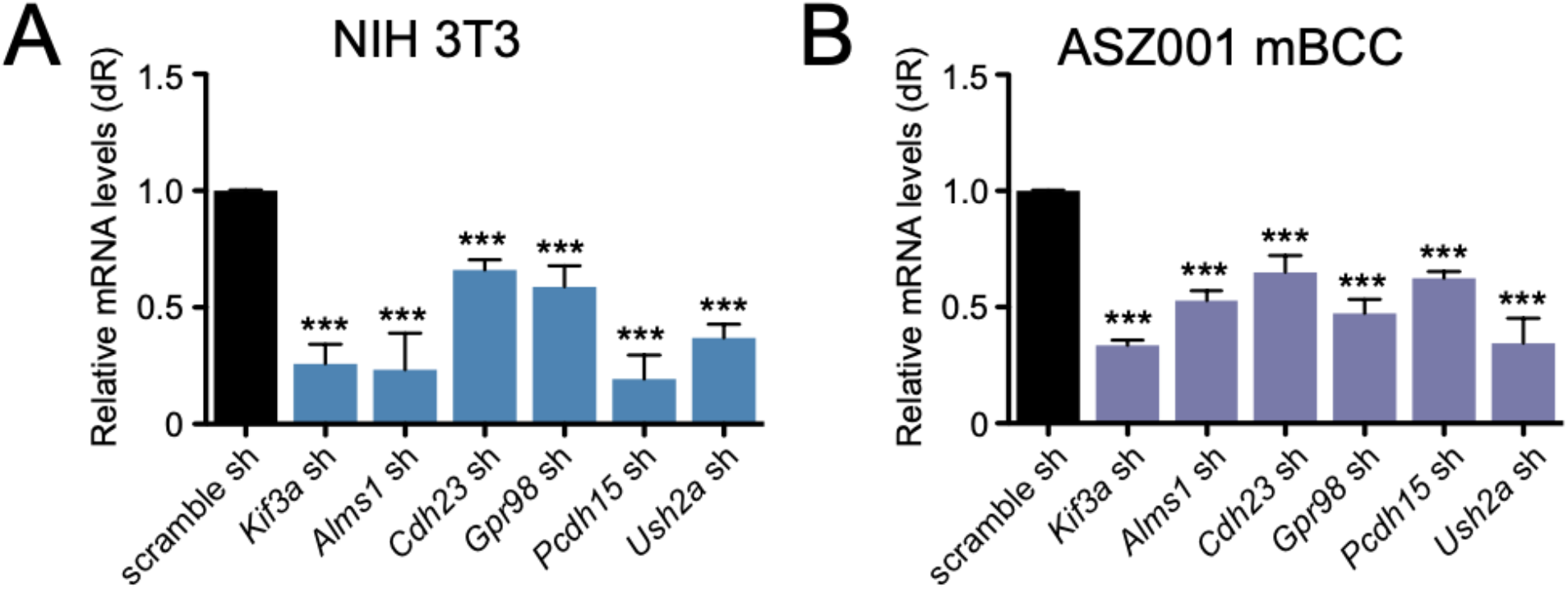
Lentiviral knockdown of Alström and Usher syndrome genes. **(A-B)** shRNA-mediated knockdown of the indicated genes in NIH3T3 cells **(A)** or ASZ001 cells **(B).** n ≥ 3. dR, delta reporter signal normalized to passive reference dye. sh, shRNA. Error bars represent SEM; significance was determined by unpaired two-tailed *t* test (***, p<0.001).

**Supplemental Figure 2.**
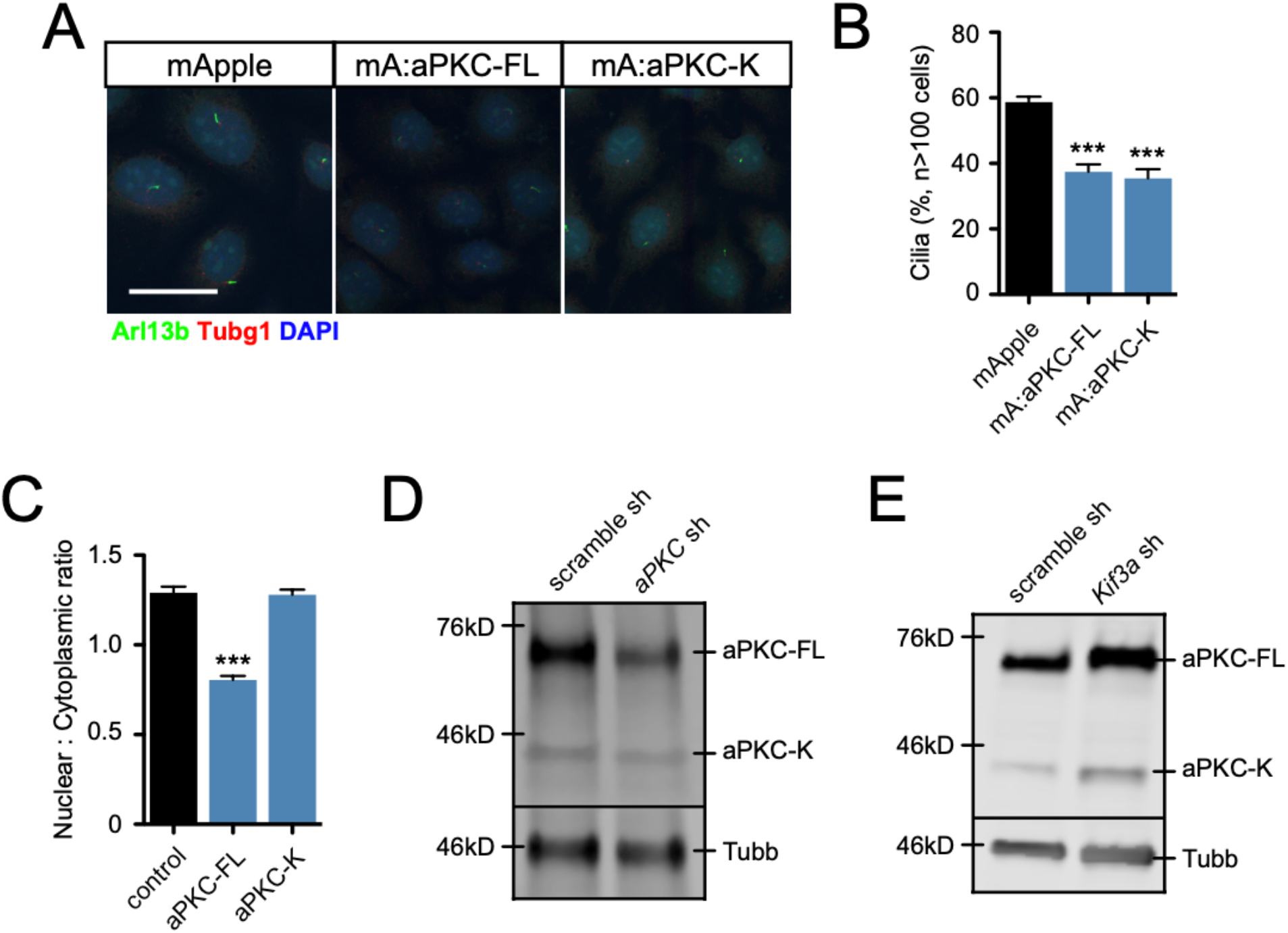
aPKC-K expression. **(A-B)** Immunofluorescence **(A)** and primary cilia quantification **(B)** of ASZ001 cells expressing the indicated constructs. Scale bar, 30 µm. mA, mApple. FL, full-length. K, kinase. %, percentage. **(C)** Nuclear to cytoplasmic ratio of the indicated constructs. **(D-E)** Western blot of *aPKC* sh-mediated knockdown **(C)** or *Kif3a* sh-mediated knockdown **(D)** using a C-terminal aPKC antibody (top panel) or Tubb antibody (bottom panel). kD, kilodalton. sh, shRNA. Error bars represent SEM; significance was determined by unpaired two-tailed *t* test (***, p<0.001).

